# Microtubule acetylation by αTAT1 is essential for touch sensation in zebrafish but dispensable for embryonic development

**DOI:** 10.1101/2025.08.07.669193

**Authors:** Samuel G. Bertrand, Daniel T. Grimes

## Abstract

Acetylation of α-tubulin at lysine 40 (α-tub^K40Ac^) is a conserved post-translational modification that occurs on the microtubule lumenal surface, but its developmental functions remain poorly defined. In zebrafish, morpholino knockdown of α-tubulin acetyltransferase 1 (αTAT1), the enzyme responsible for depositing α-tub^K40Ac^ marks, has been reported to cause severe developmental defects, whereas genetic loss-of-function studies in mice found no overt role in development. Here, we generated αTAT1 loss-of-function alleles in zebrafish and found that, in contrast to morphants, mutants are viable, fertile, and develop normally. αTAT1 mutants lack detectable α-tub^K40Ac^ in all examined tissues, indicating that no other enzyme compensates for loss of αTAT1. Both cilia and neurons normally display high levels of α-tub^K40Ac^ and despite the complete loss of this modification in αTAT1 mutants, gross cilia structure and motility were preserved, and cilia-dependent developmental processes remained intact. However, αTAT1 mutants did exhibit defects in touch responsiveness, something which could be rescued by wild-type but not catalytically inactive αTAT1. These findings demonstrate that αTAT1 is solely responsible for α-tub^K40Ac^ in zebrafish and that, while dispensable for embryonic development and ciliary function, this modification is required for normal somatosensory behavior.

## Introduction

Microtubules (MTs) are essential components of the cytoskeleton, playing key roles in processes such as cell division, cellular organization and cell-cell communication (Travis et al., 2022; Hilgendorf et al., 2024). Structurally, MTs are composed of repeating heterodimers of α- and β-tubulin, which assemble into hollow cylindrical structures with a central lumen. MTs perform their diverse roles through interactions with MT-associated proteins (MAPs). Additionally, tubulins undergo various post-translational modifications (PTMs), including acetylation, phosphorylation, glycylation, detyrosination, and glutamylation, which alter the properties of MTs and regulate their interactions with MAPs (Janke and Magiera, 2020).

One such PTM, the acetylation of the ε-amino group of lysine-40 of α-tubulin (α-tub^K40Ac^), occurs in long-lived MTs, including those in primary and motile cilia, and neuronal axons (L’Hernault and Rosenbaum, 1983; Piperno and Fuller, 1985; Cambray-Deakin and Burgoyne, 1987; LeDizet and Piperno, 1986, 1987). While most PTMs take place on the C-terminal tails of tubulin that protrude from the outer surface of the MT filament, α-tub^K40Ac^ is unique in occurring on the inner lumenal surface. This PTM is catalyzed by the highly conserved enzyme, α-tubulin acetyltransferase 1 (αTAT1), across species (Akella et al., 2010; Shida et al., 2010). α-tub^K40Ac^ is thought to increase mechanical resilience in MTs, reducing breakages and allowing MTs to persist (Eshun-Wilson et al., 2019; Portran et al., 2017; Xu et al., 2017). Alongside the enzymatic functions of αTAT1, the protein also contains nuclear import and export sequences in its disordered C-terminus. These sequences may regulate its dynamic localization and access to cytoplasmic MTs. Emerging evidence also implicates αTAT1 in nuclear functions, including regulation of gene expression and DNA damage repair (Wu et al., 2018; Ryu and Kim, 2020; Li et al., 2021; Ko et al., 2021). Thus, αTAT1 functions extend beyond MT acetylation and may include important nuclear roles.

Despite the evolutionary conservation of αTAT1 and α-tub^K40Ac^, their roles in cell function and embryonic development remain unresolved. In *Tetrahymena*, αTAT1 is not essential for survival, but loss of α-tub^K40Ac^ led to increased MT depolymerization (Gaertig et al., 1995; Akella et al., 2010). In contrast, *Toxoplasma* cells deficient in αTAT1 exhibited morphological and division defects (Varberg et al., 2016). In *C. elegans*, αTAT1 is expressed only in mechanosensory neurons and is required for touch sensation (Zhang et al., 2002; Akella et al., 2010; Shida et al., 2010; Topalidou et al., 2012). αTAT1 knockout mice were viable but showed impaired sperm motility, leading to reduced male fertility (Kalebic et al., 2013a). Additionally, αTAT1 knockout mice displayed decreased touch sensation and malformations in the dentate gyrus, a region of the hippocampus (Kalebic et al., 2013a; Kim et al., 2013).

In contrast to loss-of-function studies in mice, morpholino oligonucleotide (MO) knockdown of αTAT1 in zebrafish was reported to cause severe morphological abnormalities during development, including body axis curvature and truncation, reduced eye size, and hydrocephalus (Akella et al., 2010). Interestingly, while *atat1*-MO-injected zebrafish embryos lacked α-tub^K40Ac^ in some axons, it was detectable in other axons and in cilia, suggesting additional lysine-40 α-tub acetyltransferases may function in this organism. However, it is also possible that incomplete knockdown or off-target MO effects contributed to these phenotypes. Therefore, we sought to investigate the impact of genetic knockout of αTAT1 in zebrafish.

## Results

### αTAT1 is dispensable for zebrafish embryonic development and growth

In zebrafish, *atat1* encodes αTAT1 and produces three splice isoforms. To assess the requirement for αTAT1, we analyzed an existing *atat1* mutant line (*atat1^sa9581^*) and generated a second allele (*atat1^b1514^*) using CRISPR/Cas9 (**Fig S1A**). Both mutations truncate the protein before the acetyltransferase catalytic domain and disrupt all three isoforms (**Fig 1A**). Consistent with this, immunofluorescence using an antibody specific to the α-tub lysine-40 acetylation (α-tub^K40Ac^) post-translational modification revealed a global reduction in *atat1* mutants (**Fig 1B**). Despite this loss of α-tub^K40Ac^, both *atat1^sa9581^* and *atat1^b1514^* mutants exhibited grossly normal development, survived to adulthood, and were fertile. Moreover, because *atat1* is maternally expressed in zebrafish (Lee et al., 2013), we also generated maternal-zygotic (MZ) *atat1* mutants, which similarly developed normally (**Fig 1C** and **S1B**). These results demonstrate that αTAT1 is dispensable for zebrafish embryonic development and growth. Unless otherwise noted, all subsequent analyses were performed using MZ *atat1^sa9581^* mutants.

**Fig 1.**
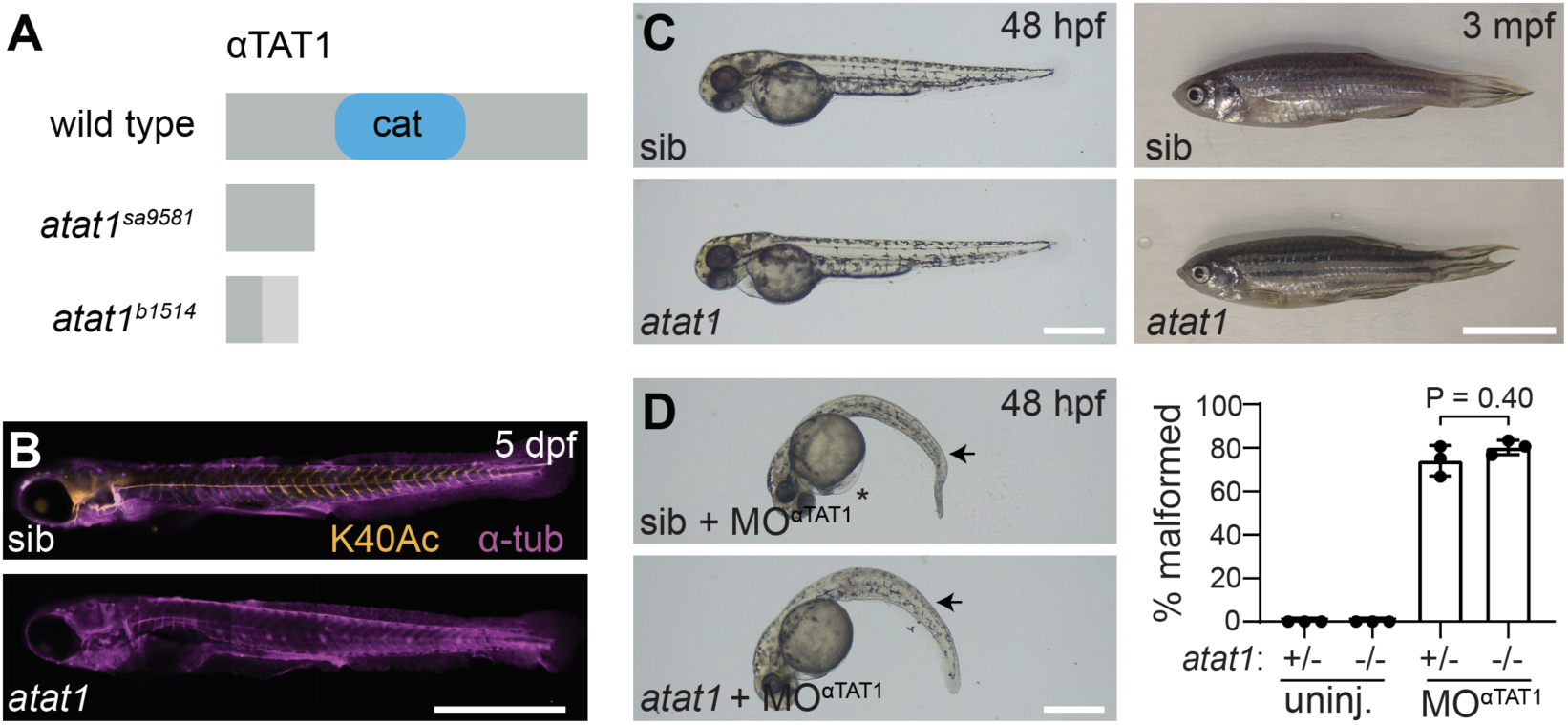
αTAT1 is not required for zebrafish development. **(A)** Schematic of the 305-amino acid αTAT1 protein, showing the N-terminal catalytic domain (cat.). The *atat1^sa9581^* allele introduces a C>T substitution at nucleotide 238, resulting in a premature stop codon. The *atat1^b1514^* allele contains a 59 bp insertion at position 83 of exon 1, also leading to a premature stop codon after 13 out-of-frame amino acids. **(B)** Immunofluorescence of 5 dpf wild-type sibling and *atat1* mutant larvae stained for α-tub^K40Ac^ and total α-tub. Scale bar, 1 mm. **(C)** (Left) Lateral views of 48 hpf wild-type sibling and *atat1* mutant embryos showing grossly normal development. Scale bar, 0.5 mm. (Right) Lateral views of 3 mpf wild-type sibling and *atat1* mutant juveniles. Scale bar, 1 cm. **(D)** (Left) Lateral views of 48 hpf wild-type sibling and *atat1* mutants embryos injected with a start-targeting morpholino (MO^αTAT1^). Asterisk indicates edema; arrows indicate body curvature. Scale bar, 0.5 mm. (Right) Percentage of larvae exhibiting malformations in uninjected and MO^αTAT1^-injected heterozygous and homozygous *atat1* mutants. *P* =0.4 (Mann-Whitney U test).

In contrast to the normal development we observed in *atat1* mutants, severe morphological defects were previously reported to be induced by morpholino oligonucleotide (MO) knockdown of αTAT1 (Akella et al., 2010). Using the same MO, called MO^αTAT1^, we found that 74.1 ± 7.0% of morphants indeed exhibited developmental malformations including curved body axes and edema (**Fig 1D**). If the phenotypes induced by MO^αTAT1^ reflect specific knockdown of αTAT1, then *atat1* mutants should be resistant to MO^αTAT1^ treatment. We therefore injected MO^αTAT1^ into *atat1* mutants and found that similar malformations were induced in *atat1* mutants, with 80.2 ± 3.3% of individuals exhibiting defects (**Fig 1D**). Thus, the MO^αTAT1^-induced phenotypes are likely to be non-specific.

### αTAT1 is essential for α-tub K40 acetylation in zebrafish cilia, but does not compromise cilia motility or function

Cilia are cell-surface organelles composed of a ring of nine microtubule (MT) doublets. Primary cilia, present on most vertebrate cells, mediate chemical signaling and mechanosensation (Hilgendorf et al., 2024), whereas motile cilia, found on specialized cell types, drive cell locomotion and generate directional fluid flows (Mitchison and Valente, 2017; Mill et al., 2023; Nachury and Mick, 2019). The axonemal MTs within cilia are highly stable and consistently modified by α-tub^K40Ac^ (L’Hernault and Rosenbaum, 1983; Piperno and Fuller, 1985; Cambray-Deakin and Burgoyne, 1987; LeDizet and Piperno, 1986, 1987). To determine whether αTAT1 is required for this modification in zebrafish and whether its loss disrupts ciliary function, we analyzed several ciliated tissues in *atat1* mutants.

We first examined cilia within the central canal (CC) of the developing spinal cord, where their beating generates cerebrospinal fluid (CSF) flow (Kramer-Zucker et al., 2005; Grimes et al., 2016). In *atat1* mutants, CC cilia completely lacked α-tub^K40Ac^ (**Fig 2A**). Nonetheless, cilia length, density, and beating frequency remained normal (**Fig 2B-E**). CC cilia-driven CSF flow is essential for assembling the Reissner fiber (RF), an extracellular glycoprotein thread composed primarily of SCO-spondin (Sspo; Reissner, 1860; Cantaut-Belarif et al., 2018). Both CSF flow and RF assembly are then required for axial straightening, the morphogenetic process by which the developing body elongates and straightens from its initial curled posture around the yolk (Bearce and Grimes, 2021). Disruption of CC cilia motility, leading to reduced CSF flow and failure to form the RF, therefore results in ventrally curved embryos (Kramer-Zucker et al., 2005). Using live imaging of an endogenously GFP-tagged Sspo (Troutwine et al., 2020), we observed robust RF assembly in *atat1* mutants (**Fig 2F**), consistent with their normal axial straightening (**Fig 1C**).

**Fig 2.**
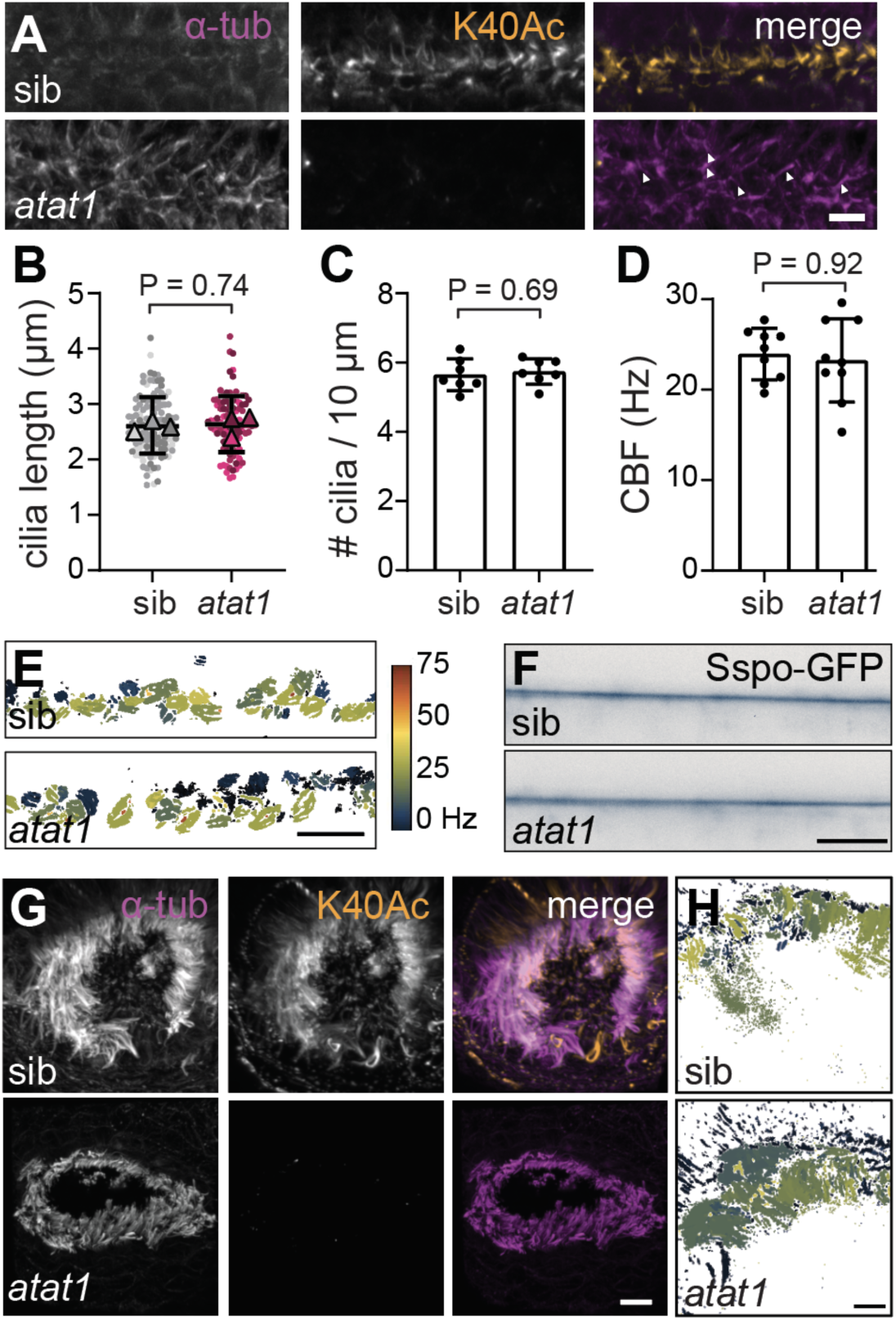
αTAT1 is essential α-tub^K40Ac^ in motile cilia. **(A)** Immunofluorescence of the central canal (CC) of 24 hpf wild-type sibling and *atat1* mutant embryos stained for α-tub^K40Ac^ and α-tub. Arrowheads point to cilia. Scale bar, 10 µm. **(B)** Superplots of CC cilia length in wild-type siblings and *atat1* mutants. Dots represent individual cilia; triangles indicate the mean for each fish (n = 3 per genotype). Bars show group mean ± standard deviation. *P* = 0.74 (unpaired *t*-test). **(C)** Cilia density in the CC, defined as the number of cilia per 10 µm, of wild-type siblings and *atat1* mutants. Bars show group mean ± standard deviation. *P* = 0.69 (unpaired *t*-test). **(D)** CC cilia beat frequency (CBF) in wild-type siblings and *atat1* mutants. Bars show group mean ± standard deviation. *P* = 0.92 (Mann-Whitney U test). **(E)** Heatmap of CBF measurements shown in (D). Scale bar, 10 µm. **(F)** Frame from live imaging of Sspo-GFP in the central canal of wild-type siblings and *atat1* mutants. Scale bar, 10 µm. **(G)** Immunofluorescence of the olfactory placode at 6 dpf in wild-type sibling and *atat1* mutant embryos stained for α-tub^K40Ac^ and α-tub. Scale bar, 5 µm. **(H)** Heatmap of CBF measurements taken from the olfactory placode, with the key as given in (E). Scale bar, 5 µm.

We made similar findings in other motile ciliated tissues, including the olfactory placode, in which motile cilia generate fluid flows that promote odor detection (Reiten et al., 2017). In *atat1* mutants, olfactory placode cilia were present and exhibited normal motility, with the average ciliary beat frequency (CBF) being 28.3 ± 4.4 Hz for *atat1* mutants and 26.7 ± 1.7 Hz for controls (*P* = 0.33, Mann-Whitney U test), despite lacking α-tub^K40Ac^ (**Fig 2G-H** and **S2B**). In the pronephros, motile cilia lining the duct were also present and motile while lacking α-tub^K40Ac^ in *atat1* mutants (**Fig S2A**).

Last, we evaluated left-right (L-R) patterning in *atat1* mutants. In zebrafish, embryonic L-R asymmetry is initiated by directional fluid flow generated by motile cilia within Kupffer’s vesicle (KV), a transient fluid-filled organ that forms in the tailbud during early somitogenesis (Essner et al., 2005; Kramer-Zucker et al., 2005; Grimes, 2019). This cilia-driven flow orients visceral organ asymmetry, and its disruption leads to randomized L-R patterning (Bisgrove et al., 2005; Essner et al., 2005; Kramer-Zucker et al., 2005; Grimes and Burdine, 2017). The earliest morphological outcome of L-R patterning is heart jogging, in which the primitive heart tube shifts leftward near the end of the first day of development (Chen et al., 1996; Grant et al., 2017). In cilia-defective mutants, this process becomes randomized (Baker et al., 2008; Smith et al., 2008; Grimes et al., 2020). *atat1* mutants, however, exhibited normal leftward heart jogging (**Fig S2C**), indicating that KV cilia remain functionally competent in the absence of αTAT1 function.

Together, these findings demonstrate that αTAT1 is required for α-tub^K40Ac^ in zebrafish motile cilia but is dispensable for ciliogenesis, cilia motility, and downstream developmental processes that rely on motile cilia function.

### αTAT1 is essential for α-tub K40 acetylation in zebrafish kinocilia but dispensable for their morphology and polarity

Having established that αTAT1 is required for α-tub^K40Ac^ in motile cilia without impairing their function, we next examined primary cilia, focusing on the mechanosensitive kinocilia of hair cells in the medial cristae of the inner ear and in lateral line neuromasts. Kinocilia in these sensory epithelia are essential for detecting mechanical stimuli and exhibit highly stereotyped morphology and orientation (Whitfield, 2020).

We first visualized hair cells in the medial cristae and quantified both kinocilia number and length. In sibling controls, kinocilia were robustly marked by α-tub^K40Ac^, but this modification was completely absent in *atat1* mutants (**Fig 3A**). Despite this loss, the number and length of kinocilia remained unchanged (**Fig 3B-C**).

**Fig 3.**
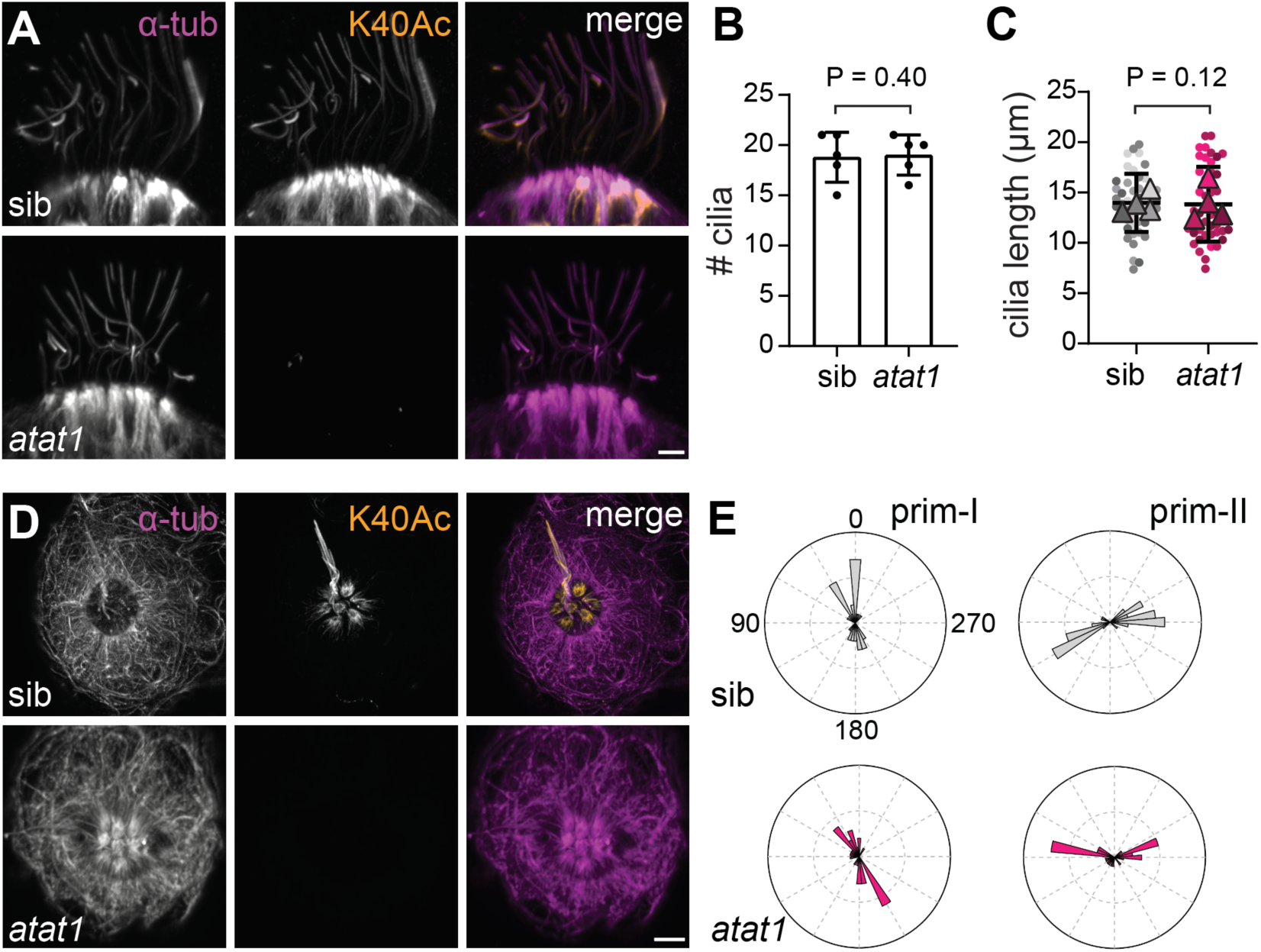
αTAT1 is essential for α-tub^K40Ac^ in kinocilia. **(A)** Immunofluorescence of medial cristae hair cells at 6 dpf in wild-type siblings and *atat1* mutants, stained for α-tub^K40Ac^ and α-tub. Scale bar, 5 µm. **(B)** Quantitation of the number of kinocilia per medial cristae in wild-type siblings and *atat1* mutants (n = 5 fish per genotype). Bars show mean ± standard deviation. *P* = 0.40 (unpaired *t-*test). **(C)** Superplots of kinocilium length in medial cristae of wild-type siblings and *atat1* mutants. Dots represent individual kinocilia; triangles indicate the mean for each fish (n = 4 per genotype). Bars show group mean ± standard deviation. *P* = 0.12 (Mann-Whitney U test). **(D)** Immunofluorescence of lateral line neuromasts in wild-type siblings and *atat1* mutants stained for α-tub^K40Ac^ and α-tub. Scale bar, 5 µm. **(E)** Rosette plots showing hair cell planar polarity in neuromasts derived from prim-I and prim-II lateral line primordia in wild-type siblings and *atat1* mutants. Hair cell orientation was measured relative to the anterior-posterior axis (0-180°), with the dorsal-ventral axis at 90-270°. Rayleigh tests of uniformity were applied to each sample: wild-type prim-I, *P* = 0.58 (n = 36 cells); wild-type prim-II, *P* = 0.90 (n = 42 cells); *atat1* prim-I, *P* = 0.08 (n = 32 cells); *atat1* prim-II, *P* = 0.19 (n = 32 cells). n = 6 fish per genotype.

Similarly, in lateral line neuromasts, α-tub^K40Ac^ was lost in the kinocilia of *atat1* mutants, but neuromast architecture appeared normal (**Fig 3D**). In these structures, pairs of hair cells exhibit opposing planar polarity, an orientation referred to as planar bipolarity (Kindt et al., 2012). This organization was preserved in *atat1* mutants (**Fig 3E**). Thus, although αTAT1 is needed for α-tub^K40Ac^ in zebrafish kinocilia, it is not required for kinocilia gross structure, length, or planar organization in sensory hair cell clusters.

### αTAT1 is required for α-tub K40 acetylation in zebrafish neurons and for touch-evoked escape responses

To assess neuronal MT acetylation, we performed immunofluorescence staining for α-tub^K40Ac^ in the trunk of 5 dpf larvae. In wild-type siblings, a prominent band of acetylated axons was visible along the spinal cord midline, corresponding to longitudinal axon tracts within the central nervous system (**Fig 4A**). In addition, segmentally arranged axons projected laterally into the somites in a chevron-like pattern, consistent with the morphology of peripheral motor nerves innervating the trunk musculature. Both the central axon bundles and peripheral projections were robustly labeled by α-tub^K40Ac^ in siblings but showed a complete loss in *atat1* mutants (**Fig 4A**), indicating αTAT1 is the enzyme responsible for α-tub^K40Ac^ in spinal and motor neurons.

**Fig 4.**
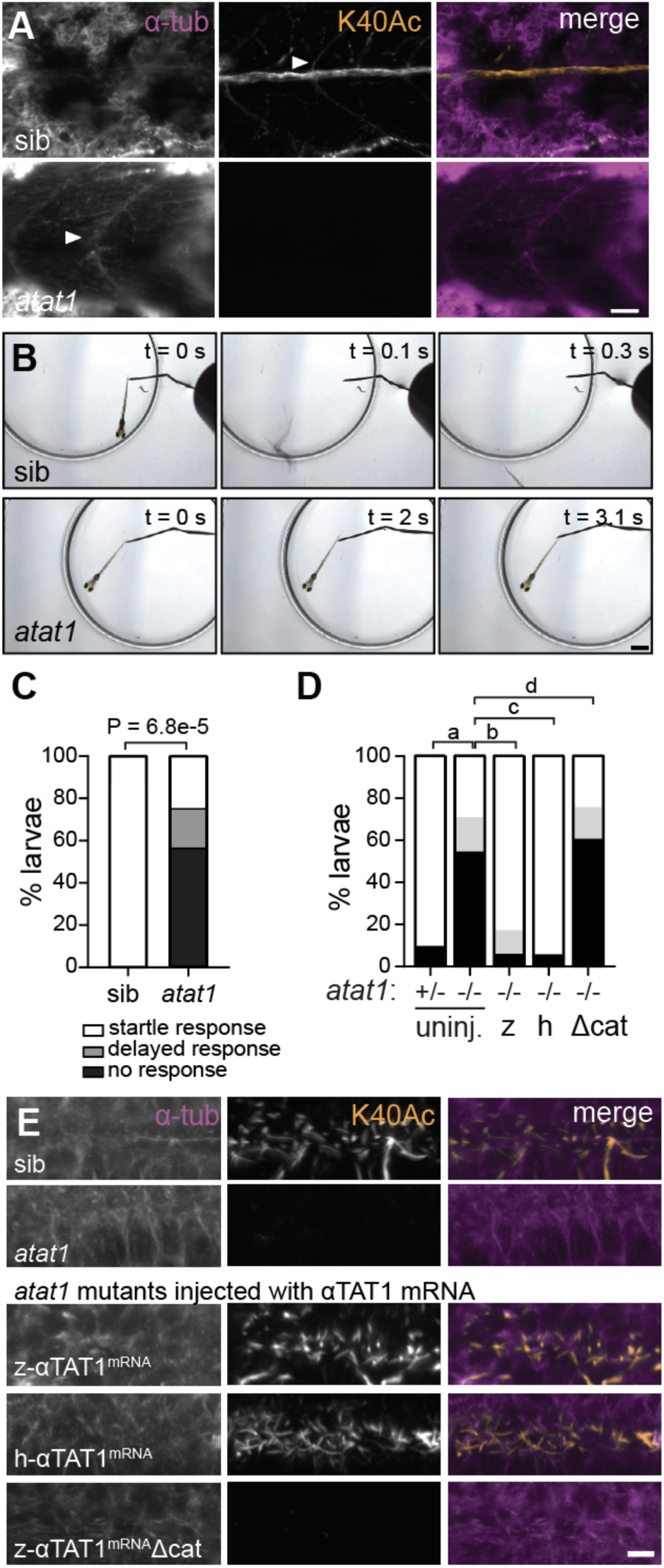
αTAT1 is essential for α-tub^K40Ac^ in neurons and touch responsiveness. **(A)** Immunofluorescence of zebrafish trunks at 6 dpf in wild-type siblings and *atat1* mutants for α-tub^K40Ac^ and α-tub. Arrowheads indicate motor neurons emanating from the spinal cord midline. Scale bar, 30 µm. **(B)** Still images from touch-evoked response assays in 6 dpf wild-type siblings and *atat1* mutants. t = 0 s marks the first touch stimulus. For *atat1* mutants, subsequent touches occurred at 2.0 s and 3.1 s. Scale bar, 1 mm. **(C)** Quantitation of touch responsiveness. A significantly greater proportion of *atat1* mutants failed to respond compared to wild-type siblings (n = 16 per group, *P* = 6.8 × 10^−5^, Chi-square test). **(D)** Quantitation of touch responsiveness for uninjected (uninj.) sibling controls (n = 22) and *atat1* mutants (n = 31), as well as mutants injected with zebrafish (z; n = 18), human (h; n = 19) and catalytically inactive (Δcat; n = 20) αTAT1 versions. Statistical comparisons using Chi-square tests: a — *P =* 4.4 × 10^−5^; b — *P* = 6.8 × 10^−4^; c — *P* = 3.5 × 10^−5^; d — *P* = 0.93. **(E)** Immunofluorescence of CC cilia in 1 dpf larvae showing α-tub^K40Ac^ and α-tub staining. Scale bar, 5 µm.

To determine whether loss of αTAT1 affects sensorimotor behavior in zebrafish, we examined touch-evoked escape responses in 5 dpf larvae. In response to a tail touch, wild-type siblings reliably initiated a stereotyped C-bend and swim burst (**Fig 4B-C**). In contrast, *atat1* mutants showed a spectrum of impaired responsiveness. The majority of mutants failed to respond even after three consecutive tail touches (n = 13/23), while a second class of mutants required two or three touches to elicit escape behavior (n = 4/23). A smaller subset responded after a single touch, similar to wild-type siblings (n = 6/23). Among *atat1* mutants that did respond to touch, both swim distance and velocity were comparable to siblings (**Fig S3A-B**). These findings indicate that, like in mice and *C. elegans*, αTAT1 is required for robust initiation of the touch-evoked escape response in zebrafish. It is possible that some responding mutants initiated movement through alternative sensory pathways, such as visual, rather than direct somatosensory detection.

The human αTAT1 protein comprises an N-terminal catalytic domain belonging to the GCN5-related N-acetyltransferase (GNAT) family and a C-terminal region that includes nuclear localization and export signals, along with an intrinsically disordered tail (Davenport et al., 2014). Zebrafish and human αTAT1 share 42% overall amino acid identity, with the highest conservation observed in the catalytic domain. Notably, all residues previously defined as critical for acetyl-CoA binding and enzymatic activity are fully conserved. Consistent with this, mRNA injection of either human or zebrafish αTAT1 robustly rescued α-tub^K40Ac^ in *atat1* mutants (**Fig 4E**), confirming that the enzymatic function is conserved across species and that the human protein can function in the zebrafish cellular context. By contrast, injection of a catalytically inactive zebrafish αTAT1 variant (Δcat), in which three residues required for acetyl-CoA binding were mutated (G128W, G130Y, and L146P; Steczkiewicz et al., 2006; Kalebic et al., 2013b), failed to restore α-tub^K40Ac^ (**Fig 4E**).

Last, we assessed whether the enzymatic activity of αTAT1 is required for its role in the touch-evoked escape response. In *C. elegans*, αTAT1 is necessary for both α-tub^K40Ac^ and touch sensitivity (Akella et al., 2010; Shida et al., 2010), but it has remained controversial whether these behavioral effects depend specifically on enzymatic activity or other aspects of αTAT1 function. Some studies have reported that catalytically inactive variants largely fail to rescue touch insensitivity (Akella et al., 2010; Shida et al., 2010), while others suggested rescue and therefore argued αTAT1’s role in touch sensitivity is independent of its enzymatic activity (Davenport et al., 2014; Topalidou et al., 2012). In mice, sensory neurons also require αTAT1 to detect mechanical stimuli, and this ability does appear to depend on αTAT1’s enzymatic deposition of α-tub^K40Ac^ marks (Morley et al., 2016). To address this question in zebrafish, we tested touch responses following mRNA rescue in *atat1* mutants. We found that while expression of either zebrafish or human αTAT1 restored the touch-evoked escape response, the catalytically inactive αTAT1 variant did not (**Fig 4D**). These results argue that enzymatic activity is essential for αTAT1’s role in mechanosensory behavior and suggest that α-tub^K40Ac^ is a conserved molecular requirement for touch sensitivity.

## Discussion

Our findings demonstrate that αTAT1 is the sole enzyme responsible for deposition of the α-tub^K40Ac^ PTM in zebrafish. Yet, its loss has limited consequences for development: zebrafish lacking both maternal and zygotic αTAT1 develop normally, reach adulthood, and are fertile, without obvious morphological defects. This resolves a discrepancy in the field, as MO-based knockdown of αTAT1 was reported to cause significant developmental abnormalities in zebrafish, including body curvature and hydrocephalus (Akella et al., 2010). Our results align with prior knockout studies in mice, in which αTAT1 deletion similarly did not cause overt developmental abnormalities (Kalebic et al., 2013a; Kim et al., 2013), and they highlight the importance of using genetic null models to assess gene function *in vivo*.

Despite the complete loss of α-tub^K40Ac^ in motile cilia, *atat1* mutants exhibited normal ciliogenesis and robust cilia motility. Downstream developmental processes that depend on motile cilia, including Reissner fiber formation, axial straightening, and L-R patterning, were similarly unaffected. Kinocilia of the medial cristae and lateral line were present and morphologically normal in *atat1* mutants, though they too lacked the α-tub^K40Ac^ PTM. We conclude that αTAT1 is largely dispensable for cilia-mediated developmental processes.

Since *in vitro* work has demonstrated that α-tub^K40Ac^ reduces the frequency of MT breakage under mechanical stress (Portran et al., 2017; Xu et al., 2017), it is possible that α-tub^K40Ac^ contributes to developmental robustness *in vivo* by protecting cilia or other MT structures under conditions of mechanical load, environmental challenge, or genetic perturbation.

One context in which αTAT1 is essential is mechanosensory behavior. We find that *atat1* mutants exhibit pronounced deficits in touch-evoked escape responses, despite being capable of swimming. This suggests the defect reflects impaired sensory detection rather than motor output. Rescue experiments demonstrate that αTAT1’s enzymatic activity is required for this function, as catalytically inactive variants fail to restore either α-tub^K40Ac^ or rescue behavioral responsiveness.

These results add to a complex and sometimes conflicting literature across species. In *C. elegans*, catalytically inactive αTAT1 has been reported to rescue touch sensation in some contexts (Davenport et al., 2014; Topalidou et al., 2012) but not others (Shida et al., 2010), while in mice, αTAT1’s catalytic activity was required for sensory neurons to detect mechanical stimuli (Morley et al., 2016). Our data support the latter model and suggest that α-tub^K40Ac^ itself, rather than a non-enzymatic function of αTAT1, is required for touch responsiveness.

Last, our results contribute to a growing view that microtubules are structurally and functionally robust even in the absence of modifications that are correlated with MT stability. Rather than being singularly required, α-tub^K40Ac^ may function as one of multiple molecular braces within a modular and partially redundant scaffold that maintains MT resilience. In this view, the absence of α-tub^K40Ac^ does not lead to the failure of MT-based systems but may render them more susceptible to mechanical strain or dysfunction under stress. Indeed, cryo-electron microscopy of *Tetrahymena* cilia revealed that α-tub^K40Ac^ subtly contributes to the structure of doublet MTs by influencing the lateral rotational angle (Yang et al., 2024). Moreover, in motile ciliary axonemes, MT inner proteins have been shown to pervade the MT lumen and enhance structural integrity (Ichikawa et al., 2017; Ma et al., 2019), raising the possibility that α-tub^K40Ac^ and MT inner proteins function together to reinforce the MT lattice. Exploring how these factors interact will be an important direction for future work, particularly in neuronal and mechanosensory systems where mechanical resilience is critical.

## Materials and Methods

### Zebrafish

AB strains of *Danio rerio* were used. Embryos from natural matings were raised at 28°C. Zebrafish lines used in this study include *atat1^sa9581^*, *atat1^b1514^* and *sspo-gfp^ut24^* (Troutwine et al., 2020). All experimental procedures were performed according to guidelines developed by the International Association for Assessment and Accreditation of Laboratory Animal Care and approved by the University of Oregon Institutional Animal Care and Use Committee.

### Generation of *atat1^b1514^* mutants

The *atat1^b1514^* line was generated using CRISPR/Cas9 mutagenesis. CRISPRscan was used to design gRNA oligos (Moreno-Mateos et al., 2015). Templates were assembled through annealing and extension of bottom strand oligo 1 (**Table 1**) using Taq Polymerase (New England Biolabs, M0273) with the following cycling parameters: 95°C for 3 min; 30 cycles of 95°C for 30 s, 45°C for 30 s, 72°C for 30 s; and a final extension of 72°C for 10 min. gRNA templates were purified using a DNA Clean & Concentrator Kit (Zymo Research, D4013) and used for *in vitro* transcription using a MEGAshortscript T7 Transcription Kit (Thermo Fisher Scientific, AM1354). gRNAs were treated with 1 µl of TURBO DNase (Thermo Fisher Scientific, AM2238) at 37°C for 15 min, and subsequently purified using an RNA Clean & Concentrator Kit (Zymo Research, R1013), aliquoted, and stored at −80°C. Single-cell embryos were injected with 200 pg of gRNAs and 100 pg of Cas9 mRNA. Mutants were screened for insertion-deletion mutations at the target site by outcrossing to wild type, extracting DNA from subsequent progeny, and performing PCR and sequencing of mutated regions. F1 families were established and outcrossed to generate F2 families. Mutations were identified by Sanger sequencing of homozygous DNA samples. *atat1^b1514^* mutants were generated using *atat1_gRNA_1* (**Table 1**), resulting in a 59 bp insertion in exon 1 that introduced 13 new residues and a premature stop codon. Genotyping was performed by PCR amplification of the exon 1 region using *atat1_exon1_1* and *atat1_exon1_2* (**Table 1**). Wild-type DNA amplification results in a 322 bp band while mutant amplification results in a 381 bp band.

**Table 1.**
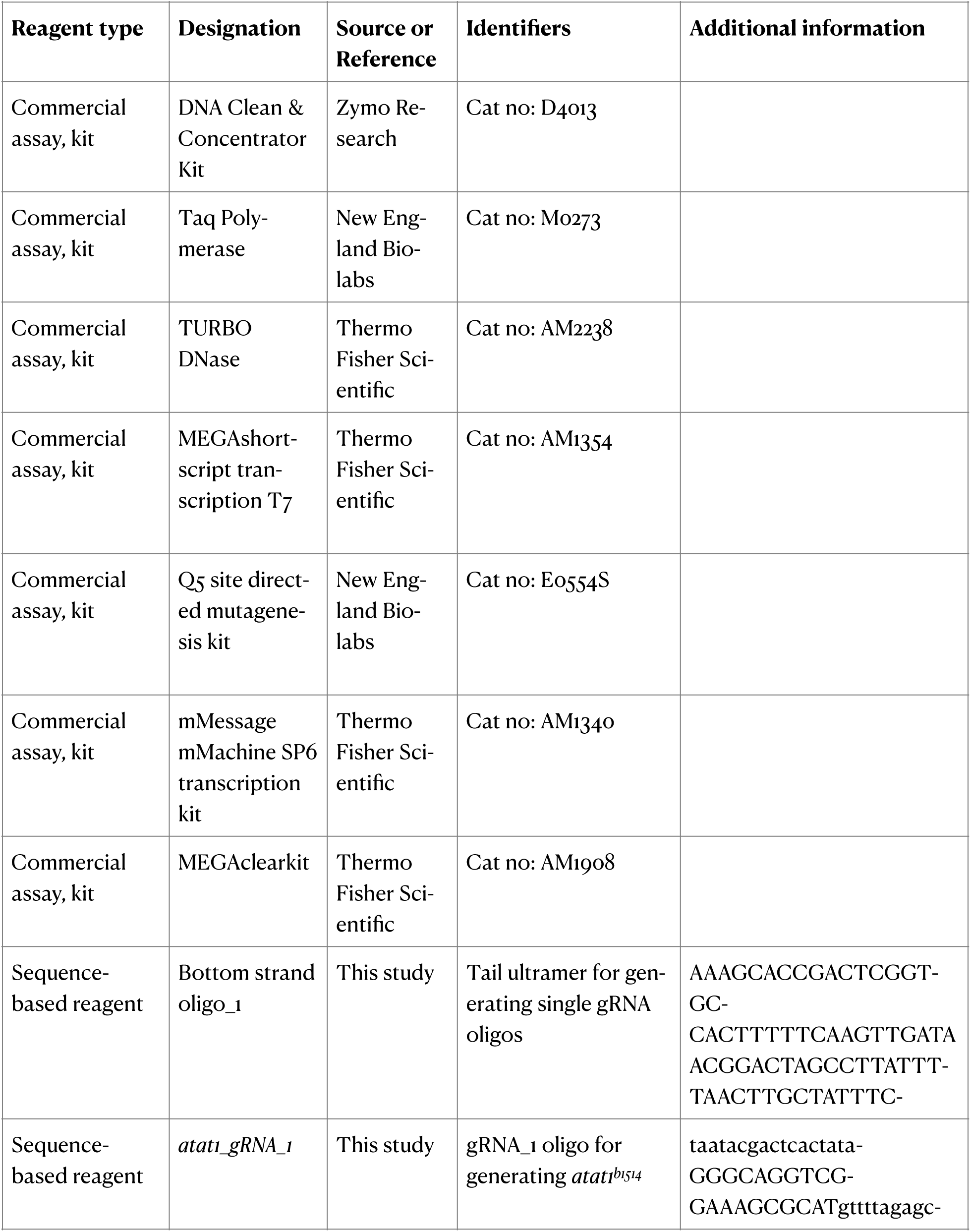
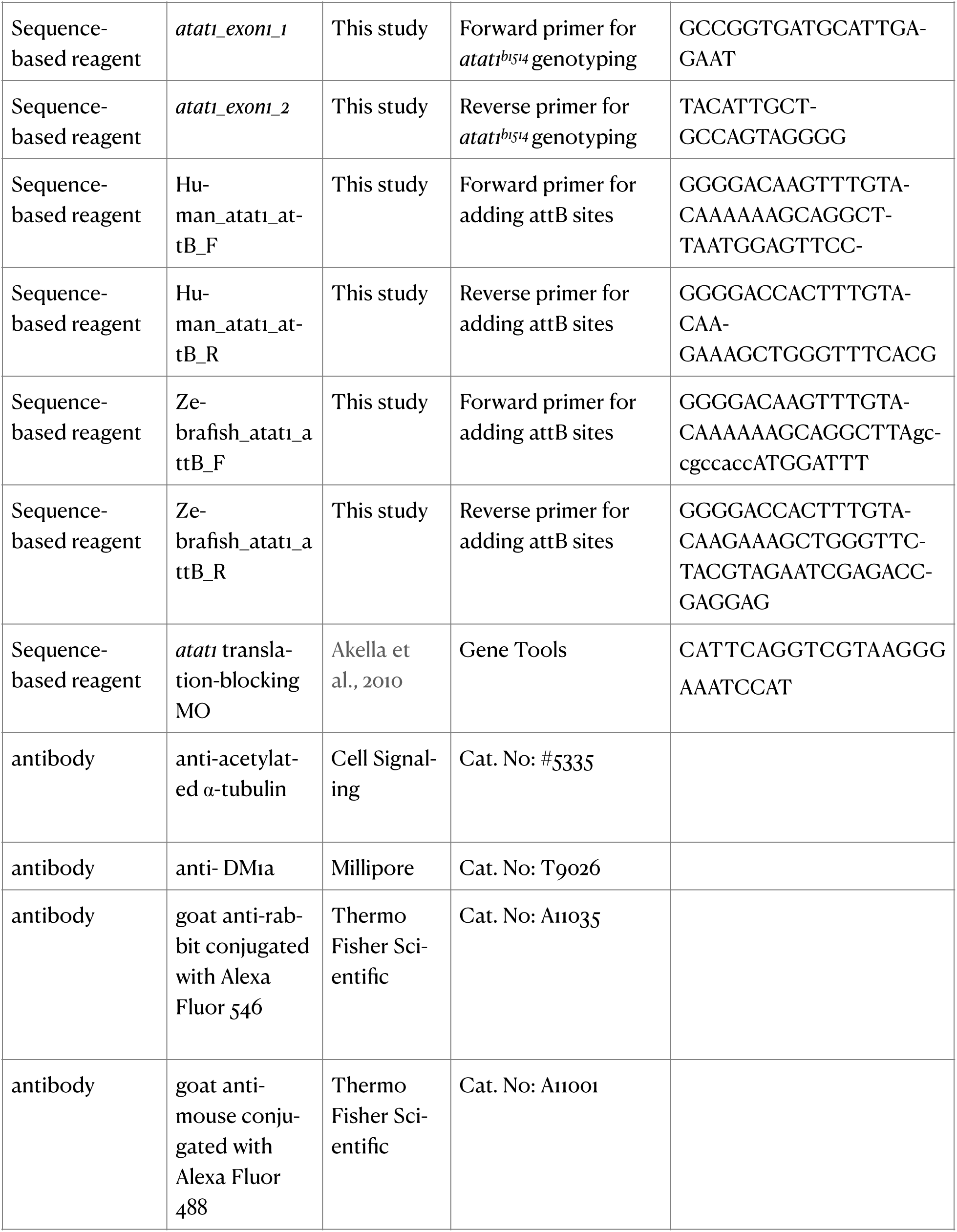
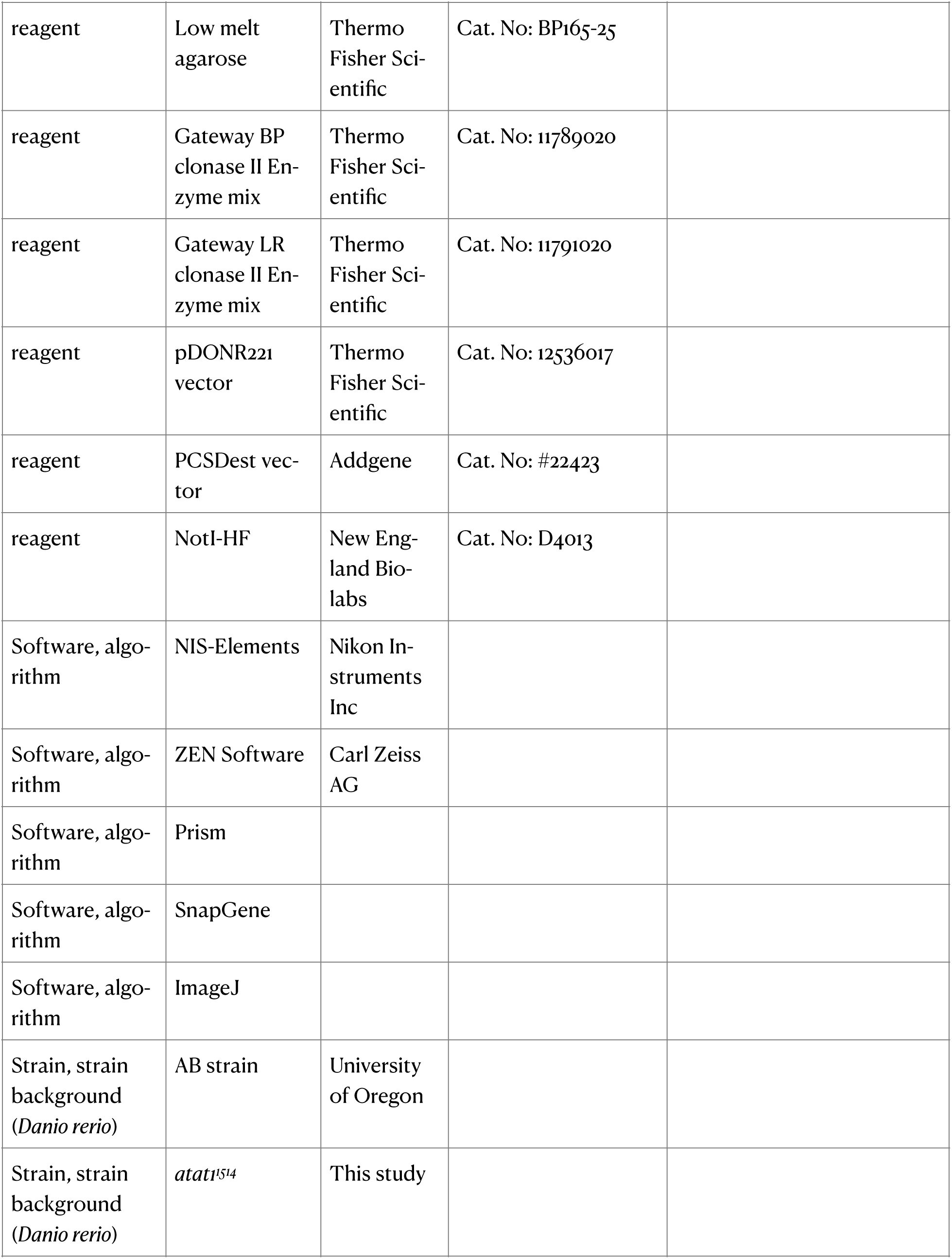
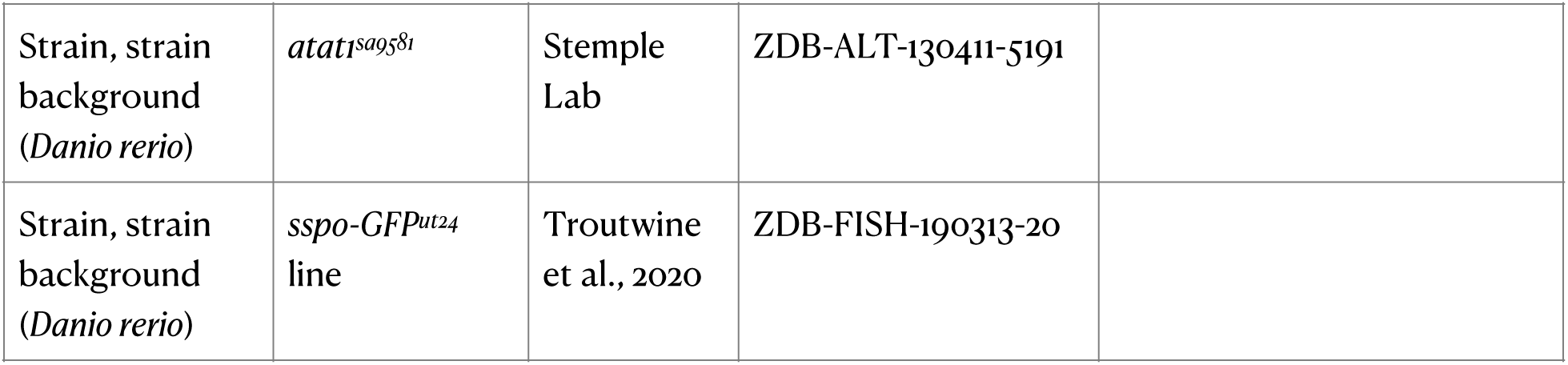
Key Resources.

### Immunofluorescence

Embryos were dechorionated and euthanized before fixation in 4% paraformaldehyde (PFA) in phosphate buffered saline (PBS) overnight at 4°C. Samples were then post-fixed in 100% methanol for either 2 hours rocking at room temperature or overnight at −20°C. Samples were rehydrated using PBS before being blocked for 2 hours in 5% normal sheep serum (NSS) and 1% dimethyl sulfoxide (DMSO) in PBST at room temperature. Primary antibody incubation in PBST containing 1% DMSO and 1% NSS occurred overnight, rocking at 4°C. Primary antibodies used included anti-acetylated α-tubulin raised in rabbit (Cell Signaling, #5335) and anti-DM1ɑ raised in mouse (Millipore, T9026). Primary antibodies were used at 1:500 dilutions. Samples were washed 5 times (>20 minutes each) in 1% NSS, 1% DMSO, and 0.1 M NaCl in PBST before being incubated in secondary antibodies at 4°C in the dark. The following secondaries were used: goat anti-rabbit conjugated with Alexa Fluor 546 (1:500 dilution, Thermo Fisher Scientific, #A11035) and goat anti-mouse conjugated with Alexa Fluor 488 (1:500 dilution, Thermo Fisher Scientific, #A11001). Samples were washed 5 times (>20 minutes each) in 1% NSS, 1% DMSO and 0.1 M NaCl in PBST. Embryos and larvae were laterally mounted on #1.5 coverslip-bottomed MatTek chambers and embedded in 0.5% low-melt agarose (Thermo Fisher Scientific, BP165-25). Laser scanning confocal microscopy was performed using PMT and GaAsP-PMT detectors, with pinhole size set to 1 Airy unit. Gain settings ranged from 650-800 (PMT). The *x/y* pixel size was 0.0852 µm. Images were deconvoluted using Huygens Professional, then processed and quantified in ImageJ. Cilia lengths were manually measured by tracing cilia using the Freehand Line tool in ImageJ. Neuromast hair cell polarity was measured by drawing vectors for each neuromast hair cell to acquire polarity angles and Rose plots were generated in R.

### Live imaging and quantitation of cilia

28-48 hpf embryos were dechorionated manually and anesthetized with tricaine before being laterally mounted on #1.5 coverslip-bottomed MatTek chambers and embedded in 0.5% low-melt agarose (Thermo Fisher Scientific, BP165-25) containing tricaine, prepared in embryo medium. For imaging of the olfactory placode, 6 dpf larvae were mounted in low melt agarose with their heads perpendicular to the bottom of MatTek chambers. Imaging was performed using a Nikon Ti2 inverted microscope equipped with a Plan Apo VC 60x WI DIC 1.2 numerical aperture objective and a pco.edge sCMOS camera. Differential interference contrast (DIC) image time series (512 x 512 pixels) were acquired at 250 frames per second for 4 seconds (1000 frames total) with an original pixel size of 0.11 µm. Image processing was performed in ImageJ (Schindelin et al., 2012). Time courses were rotated and cropped to isolate the central canal. A moving average of 55 frames was subtracted from each frame using the Subtract Moving Average function (Stowers Institute ImageJ Plugins > jay_unruh > Detrend). A Gaussian blur (s = 1.2) was then applied to all frames. For automated analysis of motile cilia frequencies, Fourier transform-based analysis was performed using the FreQ plugin (Jeong et al., 2022) to extract oscillations in pixel intensity associated with ciliary movement from processed DIC images. The following FreQ parameters were used: Recording frequency: 250 Hz; Sliding window to smooth power spectrum: 0.0; Remove background from power spectra: Yes; Percent of lowest-power pixels: 2.0; Fold SD: 1.5; Extract signal regions local SD filter: Yes, 3.0 (increase range = yes); Min accepted frequency: 5 Hz; Max accepted frequency: 75 Hz.

### Live imaging of Sspo-GFP

To assess Reissner fiber (RF) formation, 28 hpf Sspo-GFP embryos were laterally mounted for live imaging as described in the Live imaging and quantitation of cilia section. A Nikon Ti2 inverted microscope equipped with a Yokogawa W1 spinning disc, a Plan Apo VC 60x WI 1.2 NA objective, and a pco.edge sCMOS camera was used for image acquisition. Images were acquired with a 500 ms exposure every 2 seconds for 1 minute (30 frames total, 512×512 pixels). The original pixel size was 0.11 µm.

### Assessment of touch response

Blinded touch response tests were conducted at 11am in a room heated to 28 °C according to Sztal et al., 2016. In brief, a petri dish filled with embryo medium was placed on an illuminated stage and video recordings were performed using a Leica S9i stereomicroscope with an integrated 10-megapixel camera. One larva was assessed at a time. Recordings began before gently touching a tungsten needle to the tail of larvae. Larvae that failed to initiate the startle response but succeeded upon the second or third touch stimulus were recorded as exhibiting “delayed” responses. After the third stimulus, if no startle response occurred, these larvae were recorded as exhibiting no response.

### mRNA synthesis and expression

Gene fragments for zebrafish and human αTAT1 were synthesized by Twist Biosciences. attB fragments were added (**Table 1**) to each gene fragment before BP cloning (Thermo Fisher Scientific, 11789020) using a pDONR221 gateway donor vector (Thermo Fisher Scientific, 12536017). Whole plasmid sequencing was performed by Plasmidsaurus to verify entry clone generation. An LR reaction was then performed (Thermo Fisher Scientific, 11791020) using a PCSdest vector donor (Addgene, plasmid #22423) and resulting expression clones were sequenced. Site directed mutagenesis of both zebrafish and human constructs were performed with a Q5 Site-Directed Mutagenesis kit (New England Biolabs, E0554S), using primers found in **Table 1**. Expression plasmids were used for mRNA synthesis after restriction enzyme digestion with NotI-HF (New England Biolabs, D4013). Linear plasmids were purified using a DNA Clean and Concentrator kit (Zymo Research, D4013) before being used as a template for *in vitro* mRNA synthesis with a mMessage mMachine SP6 Transcription kit (Thermo Fisher Scientific, AM1340). mRNA was purified with a MEGAclear kit (Thermo Fisher Scientific, AM1908) and stored at −80°C before use. For overexpression, 80 pg of mRNA was injected at the single cell stage of development.

### Morpholino oligonucleotide injection

Morpholino (MO) knockdown of *atat1* was performed as previously described (Akella et al., 2010). A translation-blocking MO targeting the start codon of *atat1* (5′CATTCAGGTCGTAAGGGAAATCCAT-3′) was obtained from Gene Tools and injected at a concentration of 1 ng per embryo at the 1-cell stage. Injections were carried out in both wild-type AB and *atat1^sa9581^* mutant backgrounds. Embryos were scored for morphological and molecular phenotypes at 48 hpf.

## Acknowledgements

We thank Tim Mason, Judy Peirce, and the Aquatics Facility at the University of Oregon for zebrafish husbandry, as well as Adam Fries and the GC3F Biological Imaging Facility. We thank Max Horrocks, Zoe H. Irons and Elizabeth A. Bearce for discussions and experimental help.

## Author contributions

Samuel G. Bertrand, Formal analysis, Investigation, Visualization, Methodology, Writing - review and editing.

Daniel T. Grimes, Conceptualization, Supervision, Funding acquisition, Investigation, Writing - original draft, Project administration, Writing - review and editing.

## Supplementary Figures

**Fig S1.**
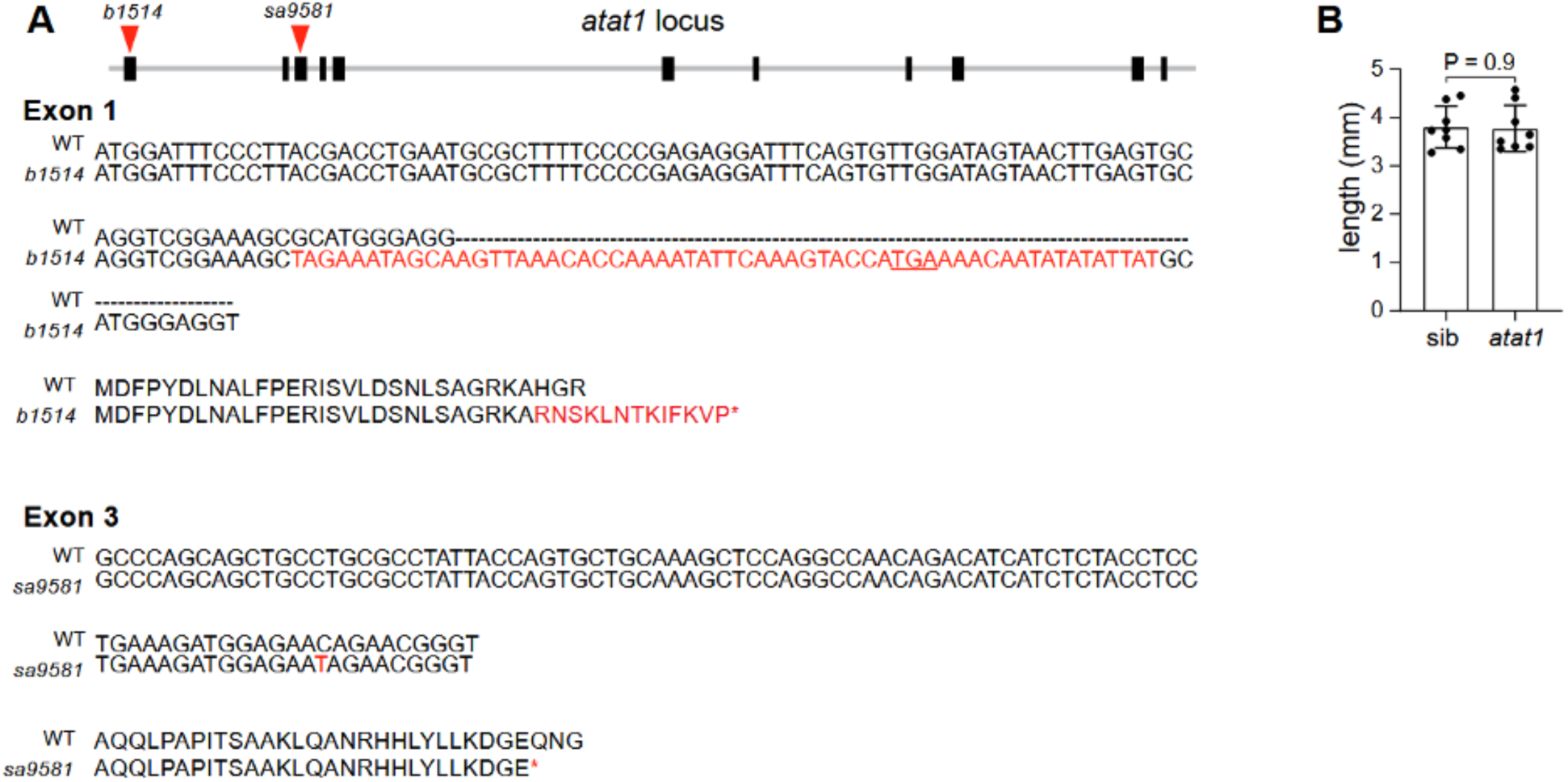
*atat1* mutant lines. **(A)** Schematic of the *atat1^b1514^* and *atat1^sa9581^* mutations. The *atat1* locus contains 11 exons. *atat1^b1514^* harbors a 59 bp insertion within exon 1 which introduces 13 additional residues before a premature stop codon (underlined). *atat1^sa9581^* harbors a single nucleotide polymorphism within exon 3 that results in a premature stop codon. **(B)** Length of wild type and *atat1* mutant adults (3 mpf). *P* = 0.9 (Unpaired *t*-test).

**Fig S2.**
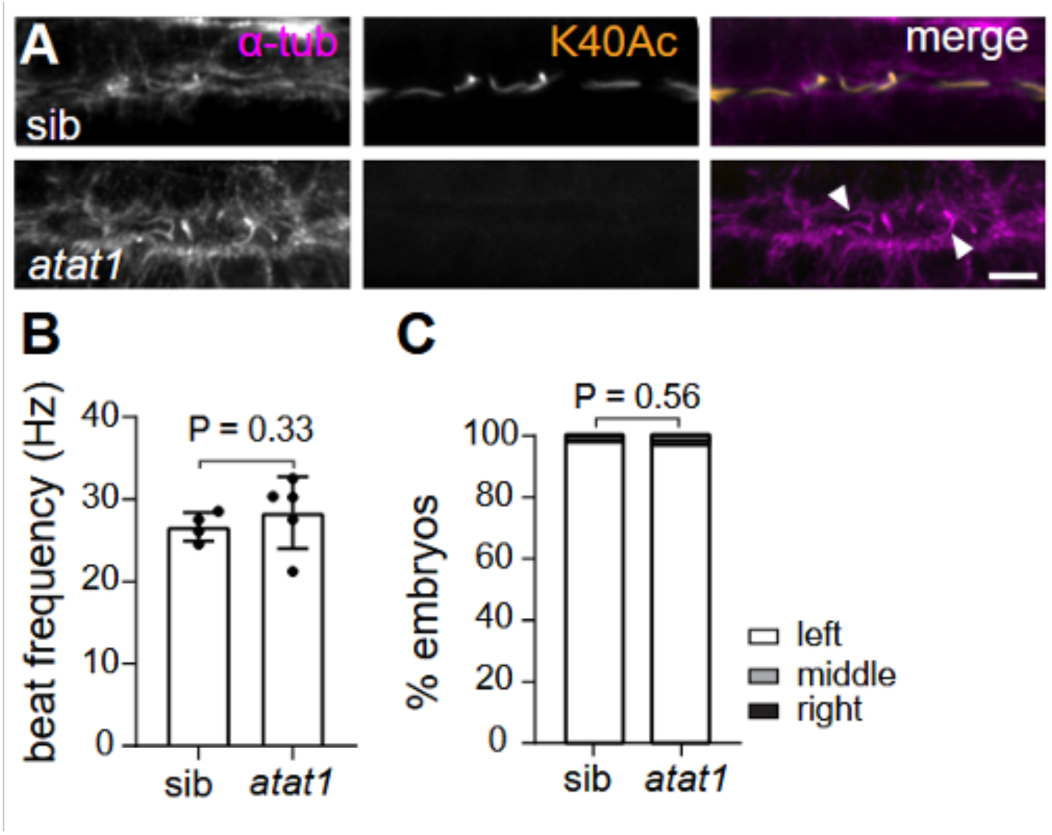
Cilia phenotyping in *atat1* mutants. **(A)** Immunofluorescence of pronephric duct cilia for wild-type and *atat1* mutants stained for α-tub^K40Ac^ and DM1ɑ (ɑ-tub). Arrows indicate cilia in *atat1* mutants. Scale bar, 5 µm. **(B)** Ciliary beat frequency measurements in the olfactory placode at 6 dpf for for wild-type siblings and *atat1* mutants. *P* = 0.33. (Mann-Whitney U test). **(C)** Percentage of embryos exhibiting left, right, or central heart jogging at 28 hpf in wild-type (n = 252) and *atat1* mutants (n = 272). *P* = 0.56 (Chi-squared test).

**Fig S3.**
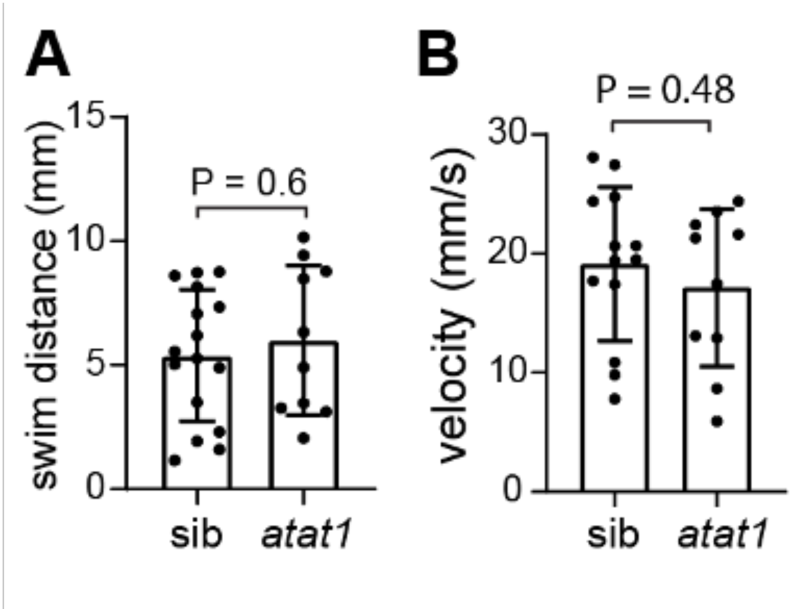
Swim distance and velocity following touch-evoked escapes. **(A)** Swim distance following touch-evoked response for wild-type siblings and *atat1* mutants categorized as exhibiting a startle or delayed response. *P* = 0.6 (Unpaired *t*-test). **(B)** Swim velocity measured in mm/s for wild-type siblings and *atat1* mutants in the startle or delayed response categories. P = 0.48 (Unpaired *t*-test).

